# A Nanobody Interaction with SARS-CoV-2 Spike Allows the Versatile Targeting of Lentivirus Vectors

**DOI:** 10.1101/2024.06.06.597774

**Authors:** Ayna Alfadhli, Timothy A. Bates, Robin Lid Barklis, CeAnn Romanaggi, Fikadu G. Tafesse, Eric Barklis

## Abstract

While investigating methods to target gene delivery vectors to specific cell types, we examined the potential of using a nanobody against the SARS-CoV-2 Spike protein receptor binding domain to direct lentivirus infection of Spike-expressing cells. Using three different approaches, we found that lentiviruses with surface-exposed nanobody domains selectively infect Spike-expressing cells. The targeting is dependent on the fusion function of Spike, and conforms to a model in which nanobody binding to the Spike protein triggers the Spike fusion machinery. The nanobody-Spike interaction also is capable of directing cell-cell fusion, and the selective infection of nanobody-expressing cells by Spike-pseudotyped lentivirus vectors. Significantly, cells infected with SARS-CoV-2 are efficiently and selectively infected by lentivirus vectors pseudotyped with a chimeric nanobody protein. Our results suggest that cells infected by any virus that forms syncytia may be targeted for gene delivery using an appropriate nanobody or virus receptor mimic. Vectors modified in this fashion may prove useful in the delivery of immunomodulators to infected foci to mitigate the effects of viral infections.

**IMPORTANCE:** We have discovered that lentiviruses decorated on their surfaces with a nanobody against the SARS-CoV-2 Spike protein selectively infect Spike-expressing cells. Infection is dependent on the specificity of the nanobody and the fusion function of the Spike protein, and conforms to a reverse fusion model, in which nanobody binding to Spike triggers the Spike fusion machinery. The nanobody-Spike interaction also can drive cell-cell fusion, and infection of nanobody-expressing cells with viruses carrying the Spike protein. Importantly, cells infected with SARS-CoV-2 are selectively infected with nanobody-decorated lentiviruses. These results suggest that cells infected by any virus that expresses an active receptor-binding fusion protein may be targeted by vectors for delivery of cargoes to mitigate infections.

## INTRODUCTION

One approach for improving the specificity of viral gene delivery vehicles is to take advantage of the surface markers of a target cell, and direct delivery by binding to such markers using nanobodies (1–3). Nanobodies offer a number of advantages for the binding of surface markers as they are small, stable, retain their binding activities when expressed extracellularly, and have an unusually long complementarity determining region (CDR) loop (4–7). We previously generated a nanobody against the receptor binding domain (RBD) of the SARS-CoV-2 Spike protein (8). This nanobody (saRBD-1), which for simplicity we will refer to as Nano, is neutralizing, and possesses a subnanomolar affinity to the SARS-CoV-2 Spike RBD (8). Because of this, we rationalized that Nano could be used to direct lentivirus vector delivery to Spike protein-expressing cells. To do so, we initially employed modified Tupaia paramyxovirus (TPMV) hemaglutinin (H) and fusion (F) proteins to pseudotype lentivirus vectors (9–10). In this system, a TPMV receptor binding-defective variant of H is equipped with C-terminal nanobody ectodomain and is incorporated along with F into lentivirus particles (9–10). Lentiviruses pseudotyped with the H-nanobody chimera (H-Nano) plus F would be predicted to infect Spike-expressing cells by virtue of H-Nano binding to Spike, and activation of the associated TPMV F protein fusion function (9–10).

Surprisingly, we found that lentiviruses pseudotyped only with H-Nano targeted gene delivery to Spike-expressing cells without the need for a TPMV F protein accessory. This implied that the H-Nano binding to Spike was enough to activate Spike’s fusion machinery and facilitate lentivirus infection. Our hypothesis was confirmed using chimeras of Nano with binding and fusion deficient variants of the HIV-1 envelope (Env) protein, and the Vesicular stomatitis (VSV) glycoprotein (G). We also demonstrated that Spike-Nano binding was sufficient to direct cell-cell fusion, and the infection of Spike-pseudotyped lentiviruses into Nano-expressing cells. We additionally found that cells infected with SARS-CoV-2 were selectively and efficiently infected with lentivirus vectors that were decorated with Nano.

Our results are consistent with observations that SARS-CoV-2-infected cells form syncytia, indicative of activated Spike proteins on infected cell surfaces (11–18). Our results also demonstrate that perturbations of host cell functions by SARS-CoV-2 infection are not sufficient to prevent lentivirus vector delivery and expression (19–21). Given that targeted lentivirus transduction to SARS-CoV-2 infected cells simply required an anti-Spike nanobody on the lentivirus surface, our findings suggest that other enveloped gene delivery vectors can be similarly targeted, either via a nanobody, or human angiotensin converting enzyme 2 (hACE2) peptide mimics (8, 22–25). More globally, our observations imply that cells infected by any syncytia-forming virus might be targeted for gene delivery in a similar fashion.

## RESULTS

### Specific gene transduction to Spike-expressing cells

We previously described an alpaca-derived nanobody, saRBD-1 against the SARS-CoV-2 Spike protein (8). This nanobody, which we will refer to as Nano, selectively binds to the Spike protein receptor binding domain (RBD; Figure 1) and neutralizes SARS-CoV-2 infection (8). We theorized that the subnanomolar affinity of the Nano-RBD interaction might be sufficient to target lentiviruses to Spike-expressing cells via pseudotyping with modified Tupaia paramyxovirus (TPMV) hemaglutinen (H) and fusion (F) proteins (9–10). As illustrated in Figure 2A, the TPMV system involves incorporation of associated, modified H and F proteins into lentivirus envelopes. The H protein variant is defective for binding to the TPMV receptor, but equipped with an alternative binding moeity, in this case Nano, yielding an H-Nano chimera. We hypothesized that binding of H-Nano to Spike would activate the fusion function of F, and result in lentivirus infection of Spike-expressing cells (Figure 2A).

**Figure 1.**
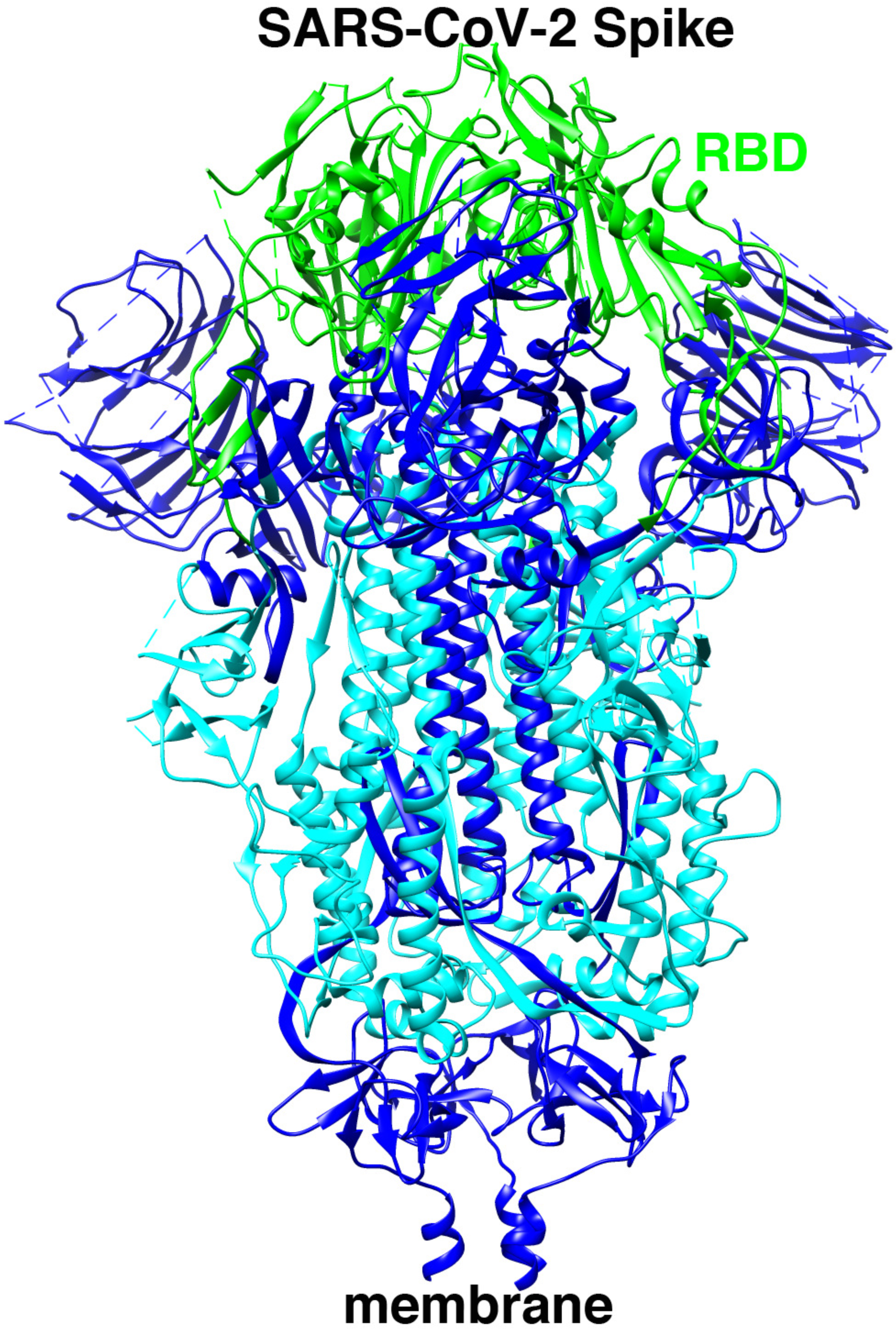
SARS-CoV-2 Spike glycoprotein. Shown is the SARS-CoV-2 Spike glycoprotein with its distal side up, and membrane-proximal side down. The receptor binding domain (RBD) in its closed conformation is illustrated in green, and residues deleted to generate a fusion-defective variant are depicted in cyan. Adapted from PDB 6VXX (58).

**Figure 2.**
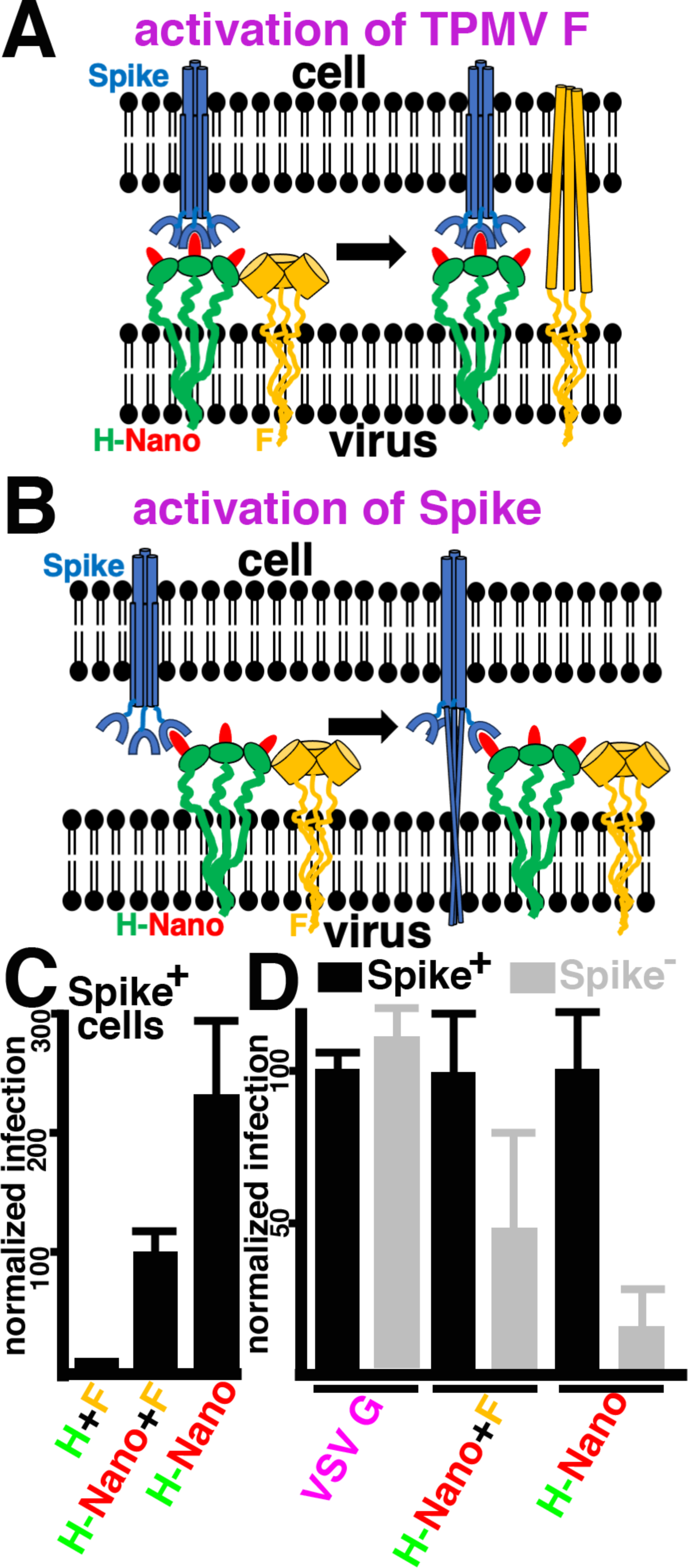
Targeting pseudotyped lentivirus vectors to SARS-CoV-2 Spike-expressing cells. **(A)** The panel depicts the cellularly expressed Spike protein in blue, with RBD regions as horseshoe shapes. The viral membrane is depicted as pseudotyped with the Tupaia paramyxovirus (TPMV) fusion (F) protein (yellow), associated with a H-Nano chimera of the binding-defective hemaglutinin (H) protein (green) and a nanobody (Nano; red) that binds to the Spike RBD. In this model, binding of the H-Nano protein to Spike activates the TPMV F protein to fuse the viral and cell membranes. **(B)** The panel depicts an alternative model, in which binding of the viral H-Nano protein to the cellular Spike protein activates Spike to fuse membranes in a TPMV F-independent fashion. **(C)** β-galactosidase (βGal) reporter lentiviruses were pseudotyped with the binding-defective TPMV H protein plus the TPMV F protein (H+F), or the H-Nano chimeric protein plus F (H-Nano + F), or the H-Nano protein alone (H-Nano). The viruses were used to infect Spike-expressing cells. Infection levels were measured via βGal activities and were normalized to the H-Nano + F levels. Averages and standard deviations are as shown. Note that the H-Nano+F and H-Nano infection levels both were significantly higher than the H+N control (P<0.001). **(D)** Spike-positive (Spike+) or Spike-negative (Spike-) cells were infected with βGal lentivirus vectors pseudotyped with the wild type (WT) Vesicular stomatitis virus glycoprotein (VSV G), or H-Nano + F, or H-Nano alone. Infection levels for each pseudotyped virus were normalized to infection of the Spike+ cells, and averages and standard deviations are as shown. Note that the difference for the H-Nano virus between Spike+ and Spike-cells was highly significant (P<0.001), while the difference for H-Nano + F was significant (P=0.002).

We tested our hypothesis on Spike-expressing cells with a β-galactosidase (βGal) transducing lentivirus vector pseudotyped either with the original F and H proteins, or with F paired with the H-Nano chimera. As illustrated in Figure 2C, the addition of H-Nano in place of H greatly increased infection of Spike-expressing cells. However, surprisingly, control viruses which included only H-Nano and not F transduced cells with even greater efficiency than the H-Nano plus F combination (Figure 2C). This immediately suggested that rather than depending on F for fusion, the interaction of H-Nano and Spike triggered the fusion function of Spike (Figure 2B). We tested the specificity for Spike-expressing cells by infection of Spike-positive (Spike+) or Spike-negative (Spike-) cells with the βGal lentivirus pseudotyped either with H-Nano plus F, H-Nano alone, or the control Vesicular stomatitis virus (VSV) glycoprotein (G). As expected, the VSV G pseudotyped lentivirus infected cells efficiently, irrespective of the presence of Spike (Figure 2D). In contrast, H-Nano plus F viruses preferentially infected Spike-expressing cells, and this disparity was increased with lentivirus vectors pseudotyped only with H-Nano (Figure 2D). These results support a model in which the presence of Nano on the lentivirus surface is sufficient to target the infection of Spike+ cells.

To confirm our model, we generated an alternative viral surface Nano chimera. To do so, the Nano coding sequence was placed between a signal sequence, and a variant of the HIV-1 transmembrane (TM, gp41) protein, which carried an N-terminal deletion that removed its fusion function. This chimera, Nano-gp41 (Figure 3A), was used to pseudotype βGal transducing lentiviruses as described above. Consistent with H-Nano results, we observed that Nano-gp41 pseudotyped viruses preferentially infected Spike-expressing cells (Figure 3B). Thus, decoration of lentivirus surfaces with the anti-Spike RBD nanobody appears to be an effective way infect Spike+ cells through activation of the Spike fusion machinery.

**Figure 3.**
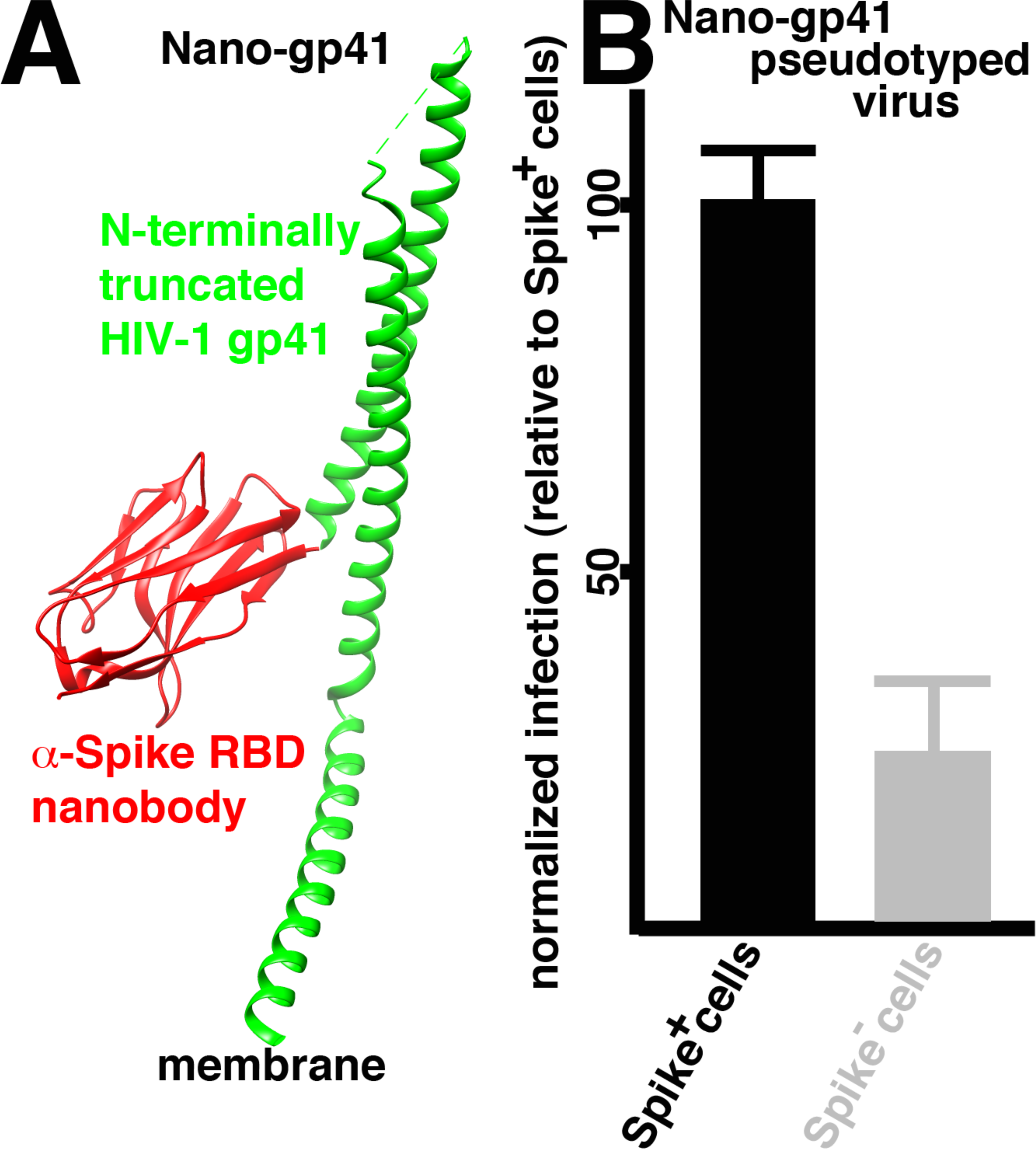
Targeting lentivirus vectors to Spike-expressing cells with a nanobody linked to the truncated HIV transmembrane (TM; gp41) protein. **(A)** The chimeric nanobody-HIV-TM protein (Nano-gp41) is composed of a N-terminal anti-Spike RBD nanobody (depicted in red by PDB 7X2L; 59) and a C-terminal, fusion-defective fragment of the HIV-1 TM protein composed of TM (gp41) residues 535-856 (depicted in green by PDB 7AEJ; 60). (B) The Nano-gp41 pseudotyped βGal lentivirus was used to infect Spike+ and Spike-cells. Infection levels were monitored by βGal activities and were normalized to values obtained for the Spike+ cells. Averages and standard deviations are as shown, and the observed difference was determined to be higly significant (P<0.001).

### Nano-Spike interactions mediate cell-cell fusion and Spike-pseudotyped lentivirus infections

Given that the Spike-Nano interaction was robust enough to fuse Spike-containing cellular membranes and Nano-expressing lentiviral membranes, we sought to determine if the interaction could facilitate the fusion of other membranes. For analysis of cell-cell fusion, we followed the experimental strategy depicted in Figure 4A. In particular, cells were separately transfected with expression vectors for Spike plus the membrane-targeted fluorescent S15-mCherry protein, or the green fluorescent protein (GFP) plus either TPMV H or H-Nano. After transfections, the cells were coincubated and monitored for red-plus green-staining syncytia. As shown in Figure 4E-G, when the control TPMV H protein was employed in the experiment, red-staining and green-staining cells were observed, but no red plus green-staining syncytia were seen. In contrast, with H-Nano, multiple red plus green-staining syncytia were evident (Figure 4 B-D). Indeed, quantitation showed that 6.8 syncytia per 100 cells were observed with H-Nano, whereas 0.24 syncytia were observed with TPMV H (Figure 4H). Thus the pairing of H-Nano and Spike in neighbor cells efficiently mediated cell-cell fusion.

**Figure 4.**
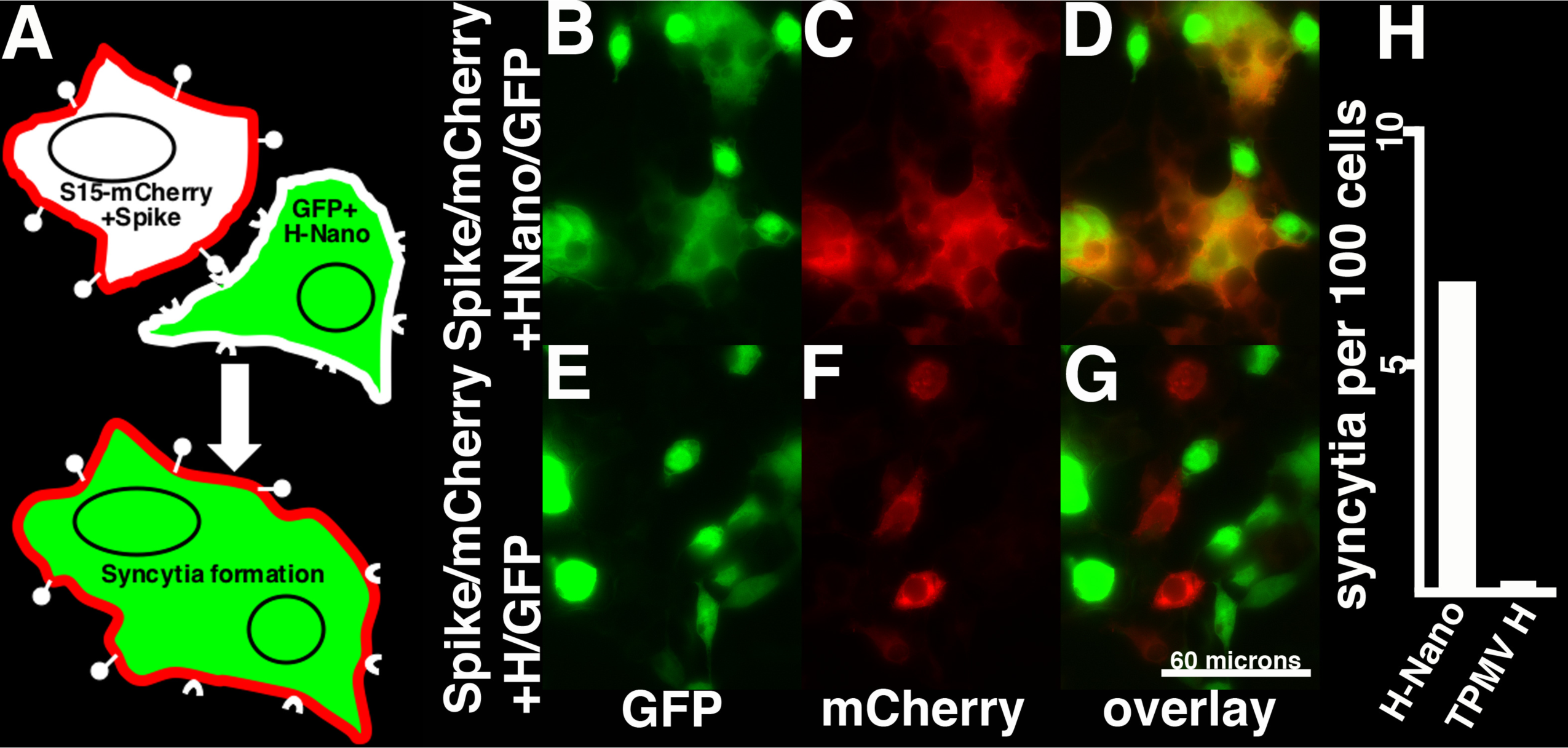
Cell-cell fusion mediated by Spike and anti-Spike nanobody interactions. **(A)** Cell-cell fusion assays were performed by coincubation of cells that had been separately transfected with expression vectors for Spike plus a membrane-targeted mCherry protein (S15-mCherry) or cytoplasm-localized GFP plus either the binding-defective TPMV H protein, or the H-Nano chimeric protein. Fusion was monitored by the appearance of GFP+ plus mCherry+ syncytia. **(B-G)** GFP (B, E), mCherry (C, F), and overlay (D, G) images of coincubations that included either the H-Nano protein (B-D) or the control TPMV H protein (E-G). Note that the 60 micron size bar in panel G pertains to all the fluorescent images (B-G). **(H)** Numbers of red plus green fluorescent syncytia for the different coincubations were normalized to every hundred cells observed in images. Note that for the H-Nano coincubation a total of 648 cells plus syncytia were counted, while 838 cells plus syncytia were counted for the control TPMV H coincubation.

We next tested whether Nano could serve as a receptor for Spike-pseudotyped lentivirus vectors. For this, Spike-pseudotyped bGal lentivirus vectors were used to infect cells expressing either H-Nano, or TPMV-H (Figure 5A). As predicted, TPMV H-expressing cells did not support infection, whereas H-Nano cells were readily transduced (Figure 5B), indicating that H-Nano can replace hACE2 as a Spike receptor.

**Figure 5.**
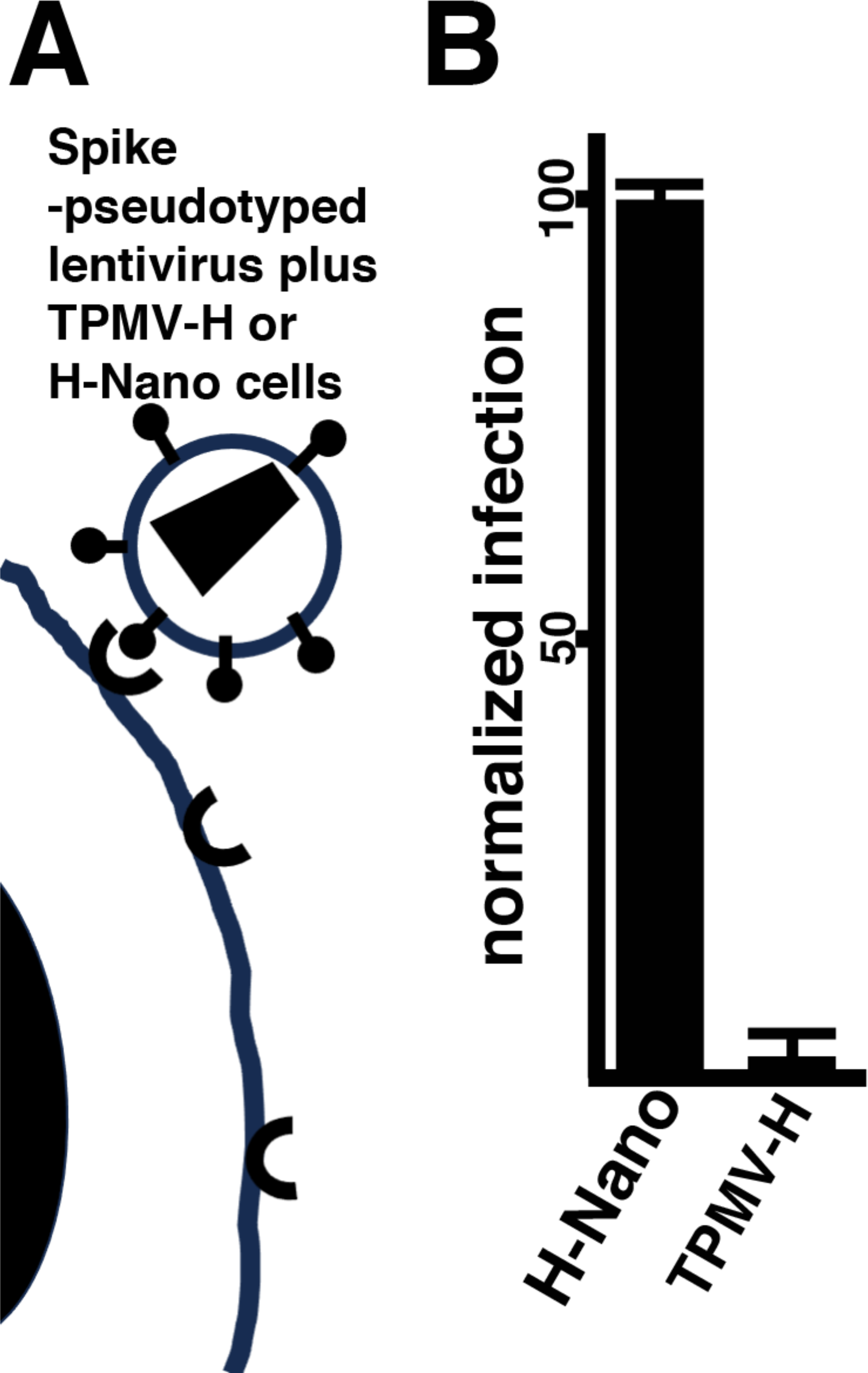
Spike-pseudotyped lentivirus infection of cells. **(A)** βGal reporter lentiviruses were pseudotyped with the SARS-CoV-2 Spike protein and used to infect cells expressing either the binding-defective TPMV H protein or the H-Nano chimeric protein that carrries the anti-Spike RBD nanobody. (B) Shown are infection levels normalized to cells expressing the H-Nano protein, with averages and standard deviations as indicated. The observed difference was calculated to be highly significant (P<0.001).

### Nano-VSV G chimeras as targeting proteins

Although our H-Nano and Nano-gp41 pseudotyped viruses capably demonstrated specific targeting of Spike-expressing cells, their infection efficiencies were considerably reduced relative to VSV G-pseudotyped lentivirus vectors. This is perhaps not surprising, since the VSV G protein is widely used for efficient lentivirus vector applications (26–27). In an effort to take advantage of this efficiency, we generated a novel G plus Nano chimera (G*+Nano). In contrast to the wild type (WT) VSV G protein (Figure 6A), the chimera was modified with Nano at the N-terminus of the G ectodomain, plus two G residue mutations (lysine 47 to glutamine [K47Q] and arginine 354 to glutamine [R354Q]) that have been shown to impair G protein binding to its major receptor, the low density lipoprotein receptor (LDL-R; Figure 6A) (28–29). We tested WT VSV G or G*+Nano pseudotyped βGal transducing lentiviruses in infections of Spike-expressing (Spike) and Spike-negative (mock) cells, and (for G*+Nano), cells expressing a SpikeΔ586-978 protein, which retains the Spike RBD in the ectodomain, but carries a deletion of residues 586-978 (see Figure 1), responsible for the Spike fusion function. As observed before (Figure 2D) viruses pseudotyped with WT VSV G showed no preference for Spike-expressing cells (Figure 6B). In contrast, G*+Nano pseudotyped viruses were highly selective for Spike-expressing cells, while viruses pseudotyped with a chimera of G* plus a negative control nanobody (G*+NanoControl) failed to transduce Spike+ cells efficiently (Figure 6B). Importantly, infection levels for G*+Nano viruses on SpikeΔ586-978-expressing cells were approximately those of Spike-minus cells, even though immunofluorescent staining patterns of WT Spike and SpikeΔ586-978 cells were similar, using an anti-Spike RBD primary antibody (Figures 6C-D). These results support an interpretation that Nano targeting of vectors to Spike-expressing cells required both a Spike-specific nanobody and the Spike fusion function. An additional observation was that infection of Spike-positive cells by G*+Nano viruses was over ten times more efficient than targeting with the H-Nano or Nano-gp41 proteins, consistent with multiple studies showing the utility of VSV G in lentivirus pseudotyping (26–27).

**Figure 6.**
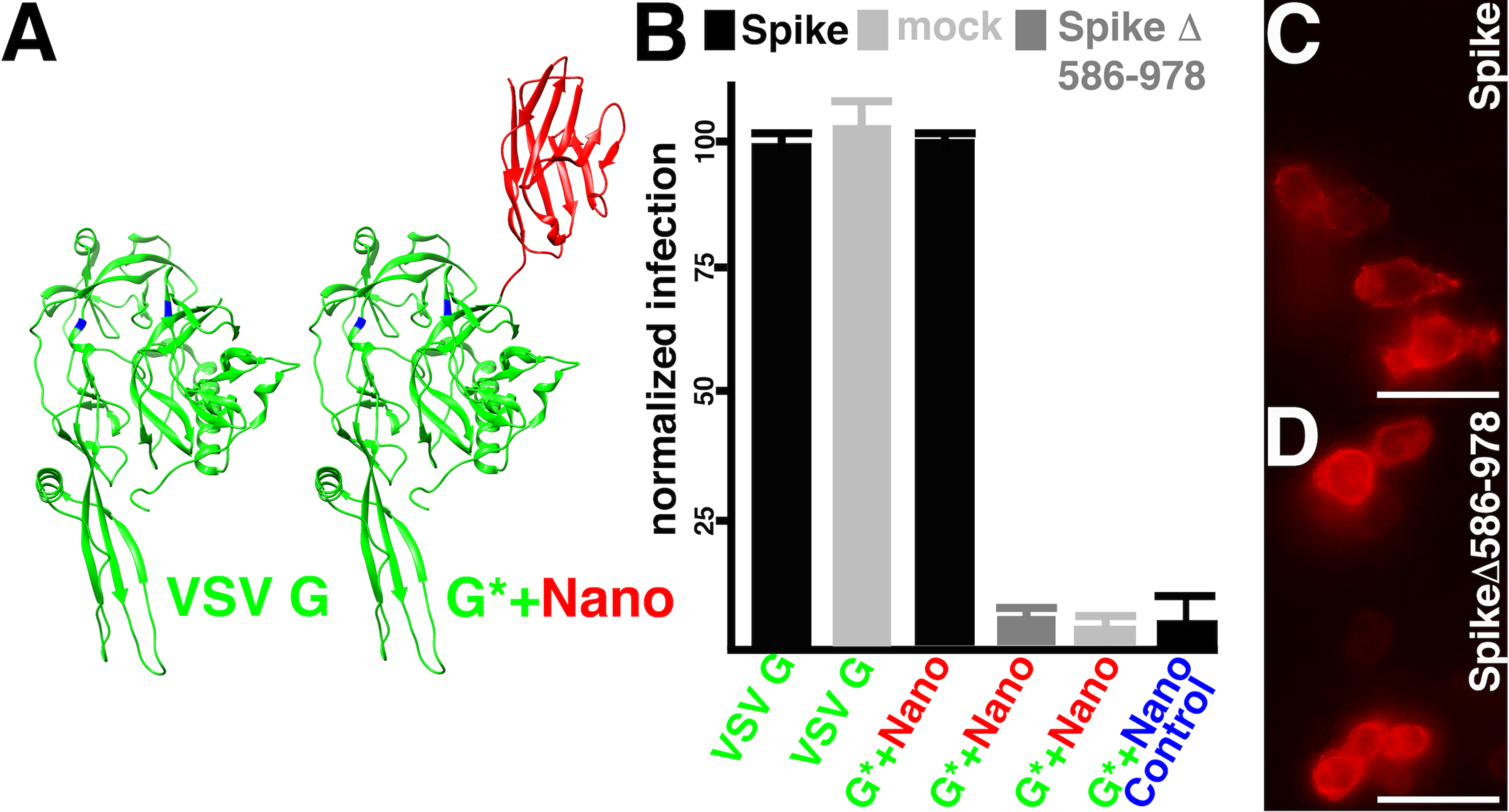
Analysis of Spike and VSV G protein variants. **(A)** VSV G protein pseudotype variants included the WT VSV G protein (VSV G, PDB 5OYL; 28), and a G*+Nano protein, which included an N-terminal anti-Spike RBD nanobody (depicted in red, and adapted from PDB 7X2L; 59) and incorporated the VSV G K47Q and R354Q mutations (in blue) that impair G protein binding to its LDL-R receptor (28). **(B)** βGal reporter lentiviruses were pseudotyped with WT VSV G or the G*+Nano protein, or a G*+NanoControl protein (with a specificity for the Epidermal Growth Factor receptor). The vectors were used to infect cells that expressed Spike (Spike+), that did not express Spike (mock), or that expressed a fusion defective variant of Spike (SpikeΔ586-978) that were deleted for the Spike residues 586-978 depicted in cyan in Figure 1. Infection levels were monitored by βGal activities and were normalized to values obtained for the VSV G-infected Spike+ cells (left two bars), or G*+Nano-infected Spike+ cells (rightmost four bars). Averages and standard deviations are as shown, and the observed differences between G*-Nano-infected Spike+ cells versus mock or SpikeΔ586-978 cells were determined to be significant (P<0.001). **(C, D)** To verify that Spike and SpikeΔ586-978 cells showed similar expression and localization patterns, cells were subjected to indirect immunofluorescent detection with a primary anti-Spike RBD antibody. Size bars indicate 30 microns.

Using G*+Nano pseudotypes, we wished to demonstrate that targeting was selective with an alternative lentivirus vector reporter. For this, we employed G and G*+Nano pseudotypes of a GFP-transducing lentivirus vector. Specifically, cells were transfected to express Spike at an efficiency of about 20% and used for infections with G- or G*+Nano-pseudotyped GFP-tranducing virus. After infections, cells were stained for Spike and GFP detection. With WT VSV G, as predicted, we observed no particular preference for infecting Spike-positive cells. This was evident in the top panels of Figure 7A, where relatively few cells stained as Spike+ and GFP+. Quantitation (Figure 7B) demonstrated that the vast majority of stained cells were only GFP+, and only half of the Spike+ cells were GFP+. In contrast, with G*+Nano, the majority of Spike+ cells were infected (Figure 7A, bottom panels), and quantitation demonstrated that three quarters of the Spike+ cells were infected, and only 5% of the infected (GFP+) cells were Spike-negative (Figure 7B).

**Figure 7.**
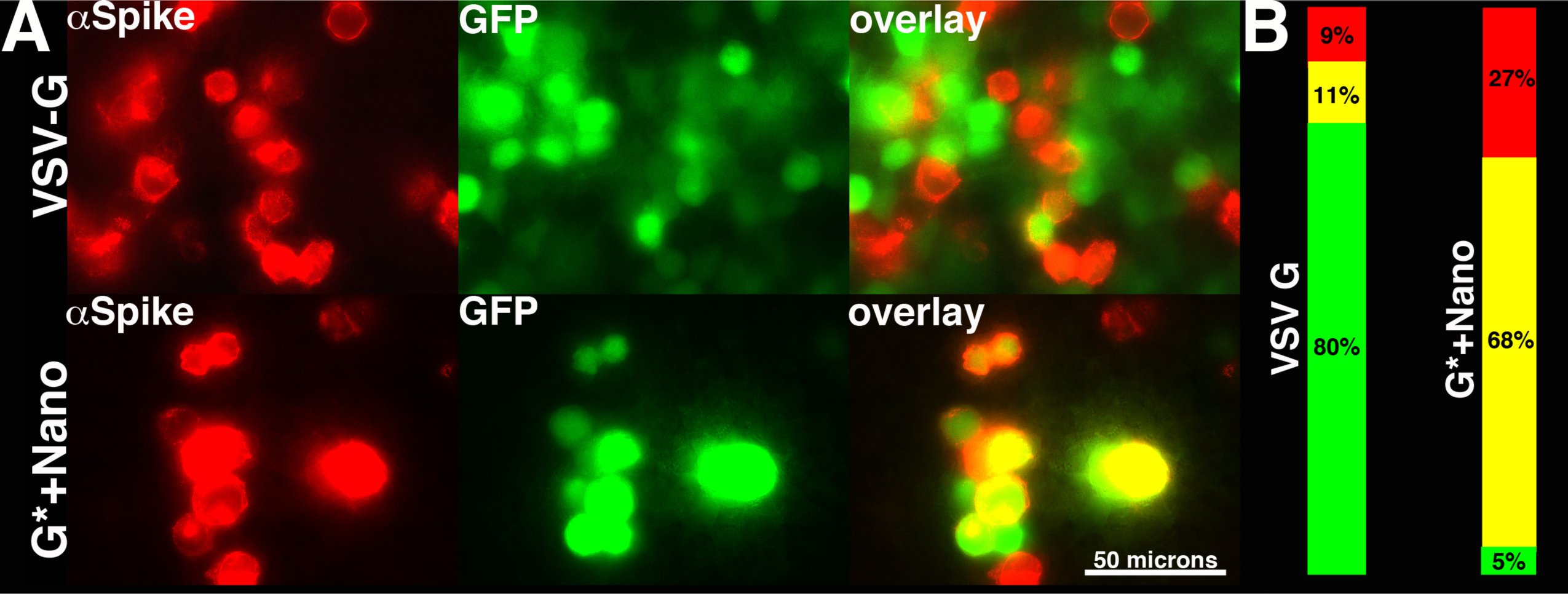
Fluorescent detection of infected cells. (**A)** GFP-transducing lentivirus vectors were pseudotyped with WT G or G*+Nano proteins and used to infect Spike-transfected cells, then stained for αSpike (red) and GFP. Note that WT VSV G transduced cells irrespective of whether they expressed Spike (top panels), but that G*+saRBD transduced cells preferentially infected Spike+ cells. (B) Quantitation GFP transduced cells staining for only GFP (green), only Spike (red), or both GFP and Spike (yellow). Note that the WT VSV G-pseudotyped vector delivered GFP to cells regardless of whether they expressed Spike, but that only 5% of cells infected with the G*+Nano virus were Spike-negative.

### Targeting lentivirus vectors to SARS-CoV-2-infected cells

While the results described above demonstrated that lentiviruses could be targeted to infect Spike-expressing cells, they did not show that SARS-CoV-2-infected cells could be targeted. For this purpose, we created a lentivirus vector that transduced expression of the human placental alkaline phosphatase (PLAP) protein. Our rationale for not using a GFP expression vector was that cytopathic effects (CPE) induced by SARS-CoV-2 infection yield autofluorescent cells that confound GFP quantitation (30–31). Our rationale for not using the βGal reporter is that infected cell lysate samples could not be removed from biosafety containment for assays without killing the βGal activity. Fortuitously, one of the features of PLAP is that it survives 60°C incubations in the presence of sodium dodecyl sulfate (SDS), which both kills endogenous alkaline phosphatase activities, and SARS-CoV-2 (32–33).

We initially tested the PLAP-transducing lentivirus in transfection experiments, where cells were transfected to express Spike, infected with WT G or G*-Nano pseudotyped PLAP vectors, and processed for PLAP assays (Figure 8A). Consistent with βGal and GFP reporters (Figures 6-7), WT G-pseudotyped PLAP viruses showed no preference for infection of Spike-expressing cells, while G*+Nano-pseudotyped viruses clearly targeted cells that had been transfected to express Spike (Figure 8C). We next followed a protocol in which Vero E6 cells first were infected with SARS-CoV-2 (strain USA-WA1/2020), and subsequently infected with pseudotyped lentivirus vectors, and then processed for PLAP detection (Figure 8B). Interestingly, the WT VSV G-pseudotyped lentivirus transduced mock-infected cells more efficiently than SARS-CoV-2 infected cells (Figure 8D), implying that SARS-CoV-2 infection either reduced the efficiency of lentivirus infection, or diminished transgene expression. Importantly, we obtained opposite results with the G*+Nano-pseudotyped lentivirus. Indeed, PLAP expression in SARS-CoV-2 infected cells was twenty times higher than in mock-infected cells (Figure 8D). Moreover, PLAP activities for SARS-CoV-2-infected cells with the G*+Nano pseudotype were more than three times higher than activities for mock-infected cells with the WT VSV G pseudotype, suggesting a highly efficient process. This efficient and selective infection of SARS-CoV-2 infected cells with G*+Nano lentivirus vectors has a number of implications that are discussed below.

**Figure 8.**
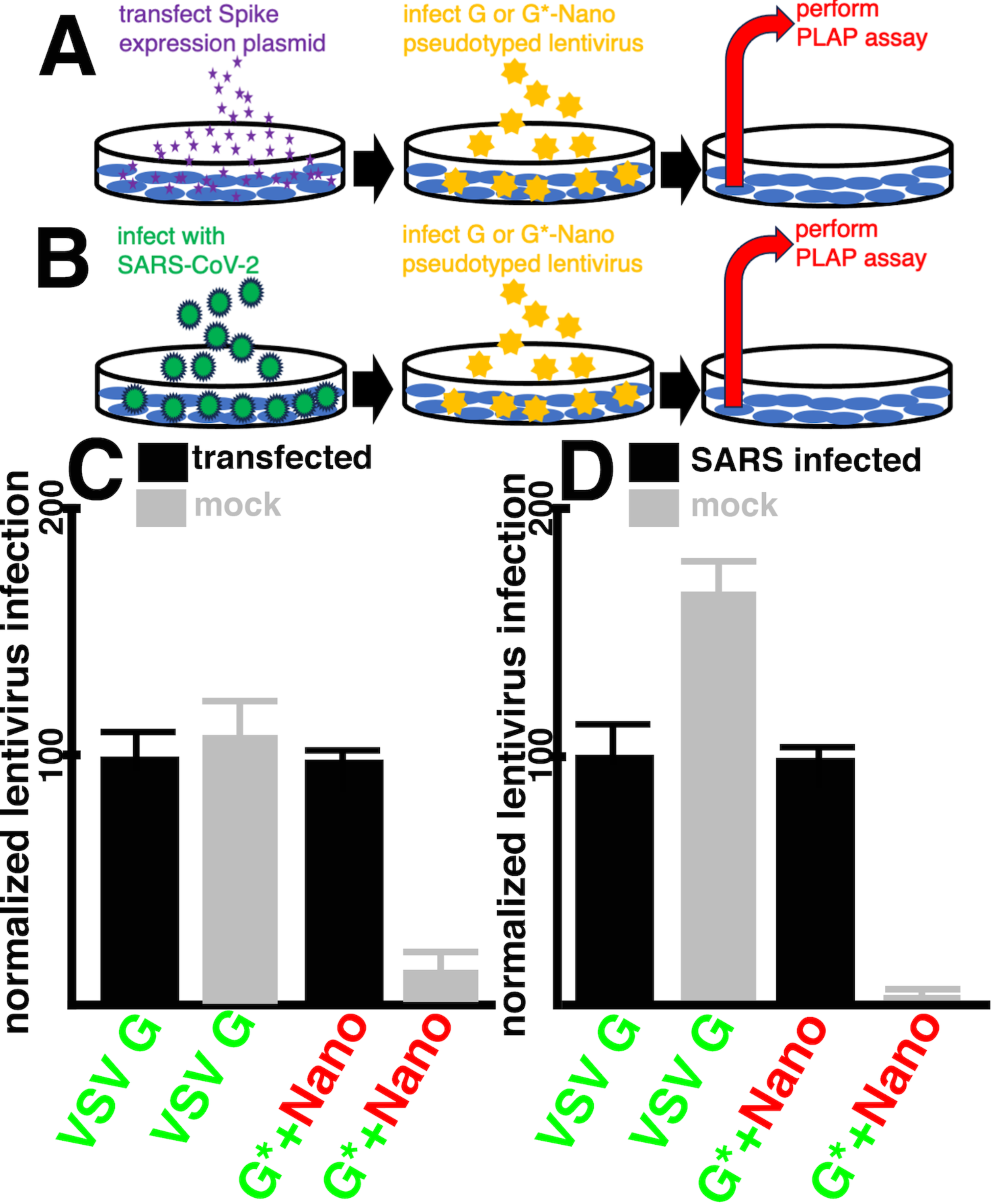
Lentiviral transduction of placental alkaline phosphatase (PLAP) to Spike-transfected or SARS-CoV-2 infected cells. **(A)** For transfection experiments, HEK293T cells were transfected with a Spike expression vector, infected at 72 h post-transfection with pseudotyped placental alkaline phosphatase (PLAP) transducing lentivirus vectors, and subjected to PLAP assays 48 h later. **(B)** For infection experiments, Vero E6 cells were infected with SARS-CoV-2 (strain USA-WA1/2020) at a multiplicity of infection (MOI) of 3, infected with pseudotyped lentivirus vectors 16 h later, and subjected to PLAP assays 28 h after that. **(C)** HEK293T cells either were mock-transfected, or transfected with a Spike expression plasmid, and subsequently infected with the indicated WT VSV G or G*+Nano pseudotyped lentivirus vectors. Infection levels for each pseudotyped virus were measured via PLAP activities, and were normalized to infection of the Spike+ cells. Averages and standard deviations are as shown. Note that the difference for the G*+Nano virus between Spike+ and Spike-cells was highly significant (P<0.001). **(D)** Vero E6 cells either were mock-infected, or infected with SARS-CoV-2, and subsequently infected with the indicated WT VSV G or G*+Nano pseudotyped lentivirus vectors. Infection levels were measured and normalized as in (C). Averages and standard deviations are as shown. Note that the difference for WT G virus between infected and mock-infected cells was highly significant (P<0.001), and the difference for the G*+Nano virus between infected and uninfected cells also was highly significant (P<0.001).

## DISCUSSION

We have demonstrated that it is possible to target the infection of SARS-CoV-2 Spike-expressing cells using lentiviruses decorated with an anti-Spike RBD nanobody (Nano) linked to modified TPMV H (H-Nano), HIV-1 TM (Nano-gp41), or VSV G (G*+Nano) proteins (Figures 2-3, 6-8). The process is dependent on the fusion function of Spike (Figure 6), and appears to follow the pathway depicted in Figure 2B, where viral Nano binding to the cellularly expressed Spike RBD activates the Spike fusion machinery. We also have shown that the Spike-Nano interaction can mediate cell-cell fusion, and infection by Spike-pseudotyped lentiviruses into Nano-expressing cells (Figures 4-5).

We observed that the efficiency of targeting was better with G*+Nano than the other variants, potentially as a consequence of higher expression, increased virion-incorporation, or better Nano presentation. In this regard, it is worthwhile to note that like WT VSV G, cellular expression of G*+Nano required the pMD.G backbone, which includes a β−globin intron, and was barely detectable in a vector that did not include an intron (pcDNA3.1). This observation is consistent with reports concerning the expression of VSV G and other of other non-codon optimized proteins (34–36). We find it interesting that G*+Nano pseudotyped lentiviruses were unable to infect SpikeΔ586-978 cells (Figure 6), because the fusion function of G* proteins carrying mutations that impair LDL-R binding (K47Q, E354Q) is reported to be active (29). Nevertheless, we also have found that G* chimeras with other nanobody sequences were similarly fusion defective. In contrast, we have found that lentivirus incorporation of G* proteins with separately expressed surface-exposed nanobodies has permitted vector targeting that was dependent on the G* fusion activity. These observations may serve to guide other vector targeting studies.

Because of the efficiency of G*+Nano pseudotyped vectors in targeting the infection of Spike-expressing cells (Figures 6, 7, 8C), it was possible to test whether such vectors could successfully target gene delivery to SARS-CoV-2-infected cells. Remarkably, targeting of SARS-CoV-2 was highly selective and highly efficient (Figure 8D). Indeed, PLAP transduction to SARS-CoV-2 infected cells was twenty times higher than to mock-infected cells, and higher than the transduction levels for the WT VSV G pseudotype.

Our results have a number of implications. An obvious one is that lentivirus vectors can deliver and express genes in SARS-CoV-2-infected cells. This was not a given, since SARS-CoV-2 possesses mechanisms for the down-regulation of non-SARS genes (19–21), and since infection could result in the activation of innate immune defenses (37–38). In this regard, we note that our lentivirus vector delivery system was able to overcome whatever Trim5α restriction was present in Vero E6 cells, even without resorting to the use of Trim-resistant Gag mutants (39–41). Another implication of our results is that it should be possible to target Spike with lentiviruses decorated with Spike-binding hACE2 fragments (24–25). We also believe that specific lentivirus delivery will not be limited to SARS-CoV-2-infected Vero E6 cells because of reports that SARS-CoV-2 infections result in syncytia formation in other cell types, and in vivo: such observations are indicative of fusion-competent Spike proteins on the surfaces of infected cells (11–18). Another consideration is that our results are likely not limited to lentiviral delivery vectors. Indeed, one can imagine that other enveloped virus-based delivery systems could be adapted to this approach, as might lipid nanoparticle-based vectors (42). We also conjecture that cells infected by other syncytia-forming viruses might be similarly targeted with appropriate nanobodies or virus receptor mimics.

It’s worthwhile asking how targeted gene delivery to virus-infected cells might be practically used to mitigate the consequences of viral infections. Indeed, an infected cell expressing enough of an envelope protein to induce syncytia formation seems unlikely to be rescued. Nevertheless, we envision that vector delivery of immunomodulators to infection foci might prove useful in the regulation of inflammation and/or the reduction of viral spread.

## MATERIALS AND METHODS

### Recombinant DNA constructs

The wild type (WT) Vesicular stomatitis virus glycoprotein (VSV G) expression plasmid (pMD.G), the Tupaia paramyxovirus (TPMV) fusion and hemaglutinin expression plasmids (pCG-TPMV-Fd32, pCG-TPMV-Hd32), the SARS-CoV-2 Spike expression plasmid (CoV2-Spike-D614G), the green fluorescent protein expression plasmid (GFP-NoPS), and the lentivirus GagPol packaging plasmid (psPAX2) all were obtained from Addgene, and were the gifts of Simon Davis, Jakob Reiser, Jennifer Doudna, and Didier Trono (10, 43–45). The plasmid that expresses membrane-bound mCherry, S15-mCherry, was obtained from the NIH AIDS Reagent Program, and was the gift of Tom Hope (46). Lentivirus transduction vectors were derived from pLVX-puro (Clontech). For β-galactosidase transduction, the small pLVX-puro XhoI-BamHI fragment was replaced with the sequence 5’ctcgagaccatggcggatcc3’, after which the βGal open reading frame from codon 9 through the termination codon from B2Bag was inserted at the BamHI site to yield pLVX-puro-Xho-ATG-βGal (47). For green fluorescent protein (GFP) transduction, the pLVX-puro-Xho-ATG-βGal XhoI-XbaI fragment was replaced by the GFP coding region from pEGFP-N1 (Clontech) to yield pLVX-puro-Xho-ATG-EGFP. For human placental alkaline phosphatase (PLAP) transduction, an EcoRI fragment containing the PLAP coding region from PinaPlap513 was first cloned into the vector pcDNA3.1 (Invitrogen) to make pcDNA3.1-PLAP513, and then the SnaBI-XbaI fragment from pcDNA3.1-PLAP513 was used to replace the SnaBI-XbaI fragment from pLVX-puro to generate pLVX-puro-PLAP513 (32).

All chimeric constructs containing the anti-Spike receptor binding domain (RBD) nanobody employed the saRBD-1 nanobody (Nano) coding region as a NotI-AscI DNA fragment where the reading frames at the two restriction sites is 5’ gcg gcc gc 3’ and 5’ g gcg cgc c 3’, and the entire Nano coding sequences is as follows: 5’ gcggccgctc aggtgcagct cgtggagtca gggggaggct tggtgcagcc tggggggtct ctgagactct cctgtgcagc ctctggaatc actttggatt attatgccat aggctggttc cgccaggccc cagggaagga gcgcgagggg gtctcatgta ttagtagtag tgatggtagc acgtcttatg cagactccgt gaagggccga ttcaccatct ccagagacac tgccaagaac acggtgtatc tgcaaatgaa cagcctgaaa cctgaggaca cggccgttta ttactgtgct tcagtacctc tgacatacta tagtggtagt taccactaca cgtgttcccc tatggggtat gacaactggg gccaggggac ccaggtcacc gtctcctcag cgcaccacag cgaagacccc tcggcgcgcc 3’ (8).

For Nano fusion to TPMV H, a type II membrane protein, the KpnI-SpeI fragment encoding the TPMV H C-terminus of pCG-TPMV-Hd32 was replaced with the sequence 5’ ggt acc ggg tct gcg gcc gcc tct aga ggg gcg cgc cac tag t 3’ to create pCG-TPMV-Hd32-NotXAscS, after which the Nano sequence was cloned into the pCG-TPMV-Hd32-NotXAscS NotI-AscI sites to yield pCG-TPMV-Hd32-Nano (H-Nano). For Nano fusion to the C-terminal fragment of the HIV-1 TM (gp41) protein, we modified the pcDNA3.1 (Invitrogen) backbone from the XhoI to ApaI site to the following sequence: 5’ ctcgagctca agcttcgaat tccgctagag ggccc 3’. Between the original pcDNA3.1 EcoRI site and the XhoI site, were inserted the following (5’ to 3’): a linker sequence of 5’ gaattccgca ttgcagagat attgtattta agtgcctagc tcgatacaat aaacgccatt tgaccattca ccacattggt gtgcacctca agct 3’, the human placental alkaline phosphatase (PLAP) signal sequence from nt 2 through the two lysines at codons 46-47 (nt 179) (48), a PstI-NotI adapter (5’ ctg cag gcg gcc gc 3’), the Nano sequence, an AscI-SmaI adapter (5’ g gcg cgc ccg ggt tcc tcc 3’), and a fragment encoding the C-terminus of the HIV-1 TM (gp41) protein from NL4-3 nt 7817-8892 (HIV-1 Env residues 535-856) (49). This construct, pcDNA3.1+XbatoEco-SARBD-gp41, expresses the Nano-gp41 protein from the pcDNA3.1 cytomegalovirus (CMV) promoter. The third Nano fusion construct, pMDG-NotAsc-VSVG-K47Q/R354Q-Nano expresses a version of the LDL-R binding-defective VSV G variant with an N-terminal Nano sequence (G*+Nano). To make this, the EcoRI site at pMD.G nt 3774 near the XbaI site was killed by addition of an A after nt 3774, retaining an ApoI site. After that, the WT VSV G coding region was replaced with an altered VSV G-encoding EcoRI cassette that carried the following changes: the non-coding sequence before the initiation codon was altered to 5’ gaattcaagc ttgtcgacag c 3’, the sequence immediately after the signal sequence was modified with a NotI-ClaI-AscI multiple cloning site (5’ aag gcg gcc gca tcg atg gcg cgc cag ttc 3’, where the first and last codons are the WT VSV G codons 17 and 18), the codons for mature VSV G residues 38-39 (full-length protein codons 54-55) were synonymously changed to 5’ ggt acc 3’ to make a KpnI site, mature residue 47 was mutated from K to Q (aag to cag), mature codons 61-62 were synonymously changed to 5’ gct agc 3’ to make a NheI site, mature codons 74-75 were synonymously changed to 5’ ggg ccc 3’ to make an ApaI site, mature codons 107-108 were synonymously changed to 5’ ccc ggg 3’ to make a SmaI site, mature codon 321 was synonymously changed to ggc to remove a KpnI site, mature codons 334-335 were synonymously changed to 5’ gat atc 3’ to create an EcoRV site, mature codon 354 was mutated from R to Q (agg to cag), mature codon 362 was synonymously mutated to ccc to remove a NdeI site, mature codons 376-377 were synonymously mutated to 5’ act agt 3’ to create an SpeI site, mature codon 388 was synonymously changed to ggc to create MscI and NcoI sites, mature codons 436-437 were synonymously changed to 5’ gag ctc’ to make a SacI site, and the coding sequence ended with 5’ aag taa ctc gag gaa ttc 3’ to create non-coding XhoI and EcoRI sites. The product of this cloning was the plasmid pMDG-NotAsc-VSVG-K47Q/R354Q, which was converted to the G*+Nano expression construct pMDG-NotAsc-VSVG-K47Q/R354Q-Nano by insertion of the Nano sequence at the NotI and AscI sites. The final expression plasmid employed, CoV2-Spike-D614GΔEcoRV, was generated by deleting the CoV2-Spike-D614 EcoRV fragment, and was used to express the SpikeΔ586-978 protein.

### Cell culture

Human embryonic kidney 293T cells (293) and Vero E6 cells were obtained from the American Type Culture Collection (ATCC) and routinely were grown in humidified 5% carbon dioxide air at 37°C in Dulbecco’s Modified Eagle’s Media (DMEM) supplemented with 10% fetal bovine sera (FBS) plus 10 mM Hepes, pH 7.3, penicillin and streptomycin (50–51). Transfections of cells were performed using polyethyleneimine (PEI) as described previously (52–53), and typically employed 24 μg total DNA for 5 million cells on 10 cm plates. For generation of WT Spike, SpikeΔ586-978, TPMV H, and H-Nano target cells, 293 cells were respectively transfected with CoV2-Spike-D614G, CoV2-Spike-D614GΔEcoRV, pCG-TPMV-Hd32, or pCG-TPMV-Hd32-Nano plasmids. For the production of pseudotyped lentivirus vectors, 293 cells typically were transfected with 8 μg each of psPAX2, a pLVX-puro reporter (pLVX-puro-βGal, pLVX-puro-EGFP, pLVX-puro-PLAP513), and one of the pseudotype expression vectors. In instances where TPMV-derived F and H proteins were tested together, transfections used 6 μg of each component (psPAX2, lentivirus vector, pCG-TPMV-Fd32, and either pCG-TPMV-Hd32 or pCG-TPMV-Hd32-Nano). For cell-cell fusion experiments, 293 cells either were transfected with 12 μg CoV2-Spike-D614G plus 12 μg S15-mCherry, or 12 μg pCG-TPMV-Hd32-Nano plus 12 μg GFP-NoPS.

Virus stocks were collected at 72 h post-transfection, filtered through 0.45 micron filters, and either used immediately, or stored at −80°C. Mock-transfected or transfected 293 targets for infections typically were split at 48 h post-transfection onto six well plates (1 million cells per well) either with or without polylysine-coated (0.01%; Sigma P4707) coverslips (Fisher 22×22-1.0). Target cells were infected 1 day later, by removal of growth media, addition of 1 ml lentivirus vector stock, incubation 4 h at 37°C, addition of 1 ml growth media, and incubation for 72 h prior to assay. Note that in experiments where immunofluorescence was performed on CoV2-Spike-D614G- or CoV2-Spike-D614GΔEcoRV-transfected target cells, the cells were split at 48 h post-transfection and subjected to immunofluorescence detection of the Spike protein the next day, as described in the immunofluorescence section. Note also that for experiments with infectious SARS-CoV-2 virus, our infection strategy was modified. In particular, Vero E6 cells were seeded at 500,000 cells per well on six well plates, and infected the next day in 2 ml of media with SARS-CoV-2 (strain USA-WA1/2020; BEI Resources NR-52281, lot 70034262) at a multiplicity of infection (MOI) of 3.0. Infections were allowed to go for 16 h, after which the virus was removed, and cells were washed three times with media. After washes, cells were infected as described above with lentivirus vectors, and infections were allowed to go for 28 h, prior to processing for assays as described below.

For our cell-cell fusion assays, separately transfected 293 cells were split onto the same coverslip (1 million cells of each transfection per coverslip) at 24 h post transfection, and co-incubated an additional 48 h, prior to processing for fluorescence imaging as described below.

### Microscopy

For direct viewing of cells expressing GFP or mCherry proteins, cells on coverslips were washed with phosphate buffered saline (PBS; 9.5 mM sodium potassium phosphate [pH 7.4], 137 mM NaCl, 2.7 mM KCl), then fixed 20 min at room temperature in 4% paraformaldehyde (PFA; Sigma) in PBS, then washed in PBS, and finally mounted onto microscope slides in Fluoro-G mounting medium. For indirect immunofluorescent detection of SARS-CoV-2 Spike either alone or in tandem with GFP detection, cells on coverslips were processed as described previously (43, 53). Briefly, cells were washed once in PBS, fixed in 4% in PBS at room temperature for 20 min, washed in PBS, permeabilized in 0.2% Triton X-100 in PBS at room temperature for 10 min, washed once, and incubated in culture medium for 10 min. Subsequently, coverslips were incubated with a 1:1000 dilution of anti-SARS-CoV-2 Spike neutralizing antibody (BEI Resources, Mouse monoclonal antibody NR-53796, 40592-MM57) in culture medium at 37 °C for 1 h, rocking every 15min. After primary antibody incubation, coverslips were washed three times in culture medium, then secondary antibody (anti-mouse Alexa-Fluor 594; Invitrogen) diluted 1:1000 in media was added, and the samples were incubated at 37 °C for 30 min, rocking once at 15 min. Following incubations, the cells were washed twice in culture medium and three times in PBS, followed by mounting onto microscope slides in Fluoro-G mounting medium.

Slides were viewed on a Zeiss AxioObserver fluorescence microscope using a 63× (Planapochromat) objective and a Zeiss filter set 10 (excitation bandpass, 450–490; beamsplitter Fourier transform, 510; emission bandpass, 515–565) for green fluorophores, or Zeiss filter set 20 (excitation bandpass, 546/12; beamsplitter Fourier transform, 560; emission bandpass, 575– 640) for red fluorophores. Images were collected with Zeiss Axiovision software with a fixed gain setting of 100 and equivalent exposure times for all samples for each fluorophore in a given experiment. For the purpose of presenting color images, gray-scale 16 bit TIF images were converted to green or red images using the Image J Image/Lookup Tables function, and then saved as 8 bit RGB images (54). For overlays of red and green images, RGB TIF images were opened in Adobe Photoshop, layered using the screen option, and flattened. For analysis of syncytia, all cells and all syncytia in multiple images were counted. Syncytia were counted as equivalent, regardless of how large they were, and at least 640 cells plus syncytia were counted for each sample. For presentation, the number of syncytia per 100 cells (including syncytia) are shown. For co-fluorescence analysis, at least 100 doubly stained (red plus green), red only, or green only fluorescent cells in multiple fields were counted, and numbers numbers of red only, green only or red plus green (yellow) were converted to percentages of the total number of cells counted.

### Enzymatic assays

β-galactosidase (βGal) and placental alkaline phosphatase (PLAP) assays were performed as described previously (32, 55). For βGal assays, media on cells in six well plate wells (35 mm diameter) were removed, and cells were scraped into 1.0 ml PBS and pelleted. Cell pellets were suspended in 150 μl PBS containing 0.1% sodium dodecyl sulfate (SDS), vortexed, supplemented with 600 μl PM-2 buffer (33 mM NaH2PO4, 66 mM Na2HPO4, 2 mM MgSO4, 0.1 mM MnCl2, 40 mM β-mercaptoethanol [BME]), vortexed, supplemented with 150 μl 4 mg/ml 2-nitrophenyl-β-d-galactopyranoside (ONPG) in PM-2 buffer and incubated at 37°C. Reactions were stopped by addition of 375 μl 1 M Na2CO3 and flash freezing on dry ice powder. Samples then were thawed, and 420 nm light absorbances were read spectrophotometrically to calculate β-galactosidase activities (1 unit = 1 nMole ONPG hydrolyzed per minute = 420 nm absorbance x 285/minutes of incubation) as a measure of infectivity (55–57).

For PLAP assays, cells were washed, scraped into PBS and pelleted as described above. Cells from each well then were suspended in 100 μl cell salts buffer (50 mM Tris pH 7.4, 150 mM NaCl, 1 mM MgCl2), supplemented with 10 μl 1% SDS, vortexed and incubated at 60°C for 60 min, which both inactivates endogenous alkaline phosphatases, and kills SARS-CoV-2 (32–33). After the heat inactivation step, samples were cooled to room temperature, supplemented with 900 μl alkaline phosphatase buffer (100 mM Tris pH 9.5, 100 mM NaCl, 5 mM MgCl2) containing 1 mg/ml para-nitrophenylphosphate, and incubated at 37°C. Reactions were stopped by addition of 100 ul 1 M NaOH and frozen. Samples then were thawed, and 420 nm light absorbances were read spectrophotometrically to calculate PLAP activity (1 milliunit=OD420 absorbance increase of 0.001 per minute of reaction). Enzyme activities were taken as proportionate to infectivity, and for standardization, values were normalized to averages of Spike-expressing (or SARS-CoV-2-infected) cells. To calculate significance values, means and standard deviations from multiple experiments were determined and converted to Z values, which were used to calculate probabilities.

## ACKNOWLEDGMENTS

We are grateful to Addgene, the NIH AIDS Reagent Program, and Simon Davis, Jakob Reiser, Jennifer Doudna, Didier Trono, and Tom Hope for making recombinant DNA constructs available. EB and FT also are thankful for funding from the National Institutes of Health through grants RO1 AI152579 (EB) and RO1 AI141549 (FT).

## Notes

### Competing Interest Statement

The authors have declared no competing interest.

## REFERENCES

1. Zhou Q, Buchholz CJ. Cell type specific gene delivery by lentiviral vectors: New options in immunotherapy. Oncoimmunology. 2013 Jan 1;2(1):e22566. doi: 10.4161/onci.22566. PMID: 23483777; PMCID: PMC3583925.

2. Frank AM, Buchholz CJ. Surface-Engineered Lentiviral Vectors for Selective Gene Transfer into Subtypes of Lymphocytes. Mol Ther Methods Clin Dev. 2018 Oct 17;12:19–31. doi: 10.1016/j.omtm.2018.10.006. PMID: 30417026; PMCID: PMC6216101.

3. Strebinger D, Frangieh CJ, Friedrich MJ, Faure G, Macrae RK, Zhang F. Cell type-specific delivery by modular envelope design. Nat Commun. 2023 Aug 23;14(1):5141. doi: 10.1038/s41467-023-40788-8. PMID: 37612276; PMCID: PMC10447438.

4. Hamers-Casterman C, Atarhouch T, Muyldermans S, Robinson G, Hamers C, Songa EB, Bendahman N, Hamers R. Naturally occurring antibodies devoid of light chains. Nature. 1993 Jun 3;363(6428):446-8. doi: 10.1038/363446a0. PMID: 8502296.

5. Beghein E, Gettemans J. Nanobody Technology: A Versatile Toolkit for Microscopic Imaging, Protein-Protein Interaction Analysis, and Protein Function Exploration. Front Immunol. 2017 Jul 4;8:771. doi: 10.3389/fimmu.2017.00771. PMID: 28725224; PMCID: PMC5495861.

6. Muyldermans S. Applications of Nanobodies. Annu Rev Anim Biosci. 2021 Feb 16;9:401–421. doi: 10.1146/annurev-animal-021419-083831. Epub 2020 Nov 24. PMID: 33233943.

7. Jin BK, Odongo S, Radwanska M, Magez S. Nanobodies: A Review of Generation, Diagnostics and Therapeutics. Int J Mol Sci. 2023 Mar 22;24(6):5994. doi: 10.3390/ijms24065994. PMID: 36983063; PMCID: PMC10057852.

8. Weinstein JB, Bates TA, Leier HC, McBride SK, Barklis E, Tafesse FG. A potent alpaca-derived nanobody that neutralizes SARS-CoV-2 variants. iScience. 2022 Mar 18;25(3):103960. doi: 10.1016/j.isci.2022.103960. Epub 2022 Feb 22. PMID: 35224467; PMCID: PMC8863326.

9. Enkirch T, Kneissl S, Hoyler B, Ungerechts G, Stremmel W, Buchholz CJ, Springfeld C. Targeted lentiviral vectors pseudotyped with the Tupaia paramyxovirus glycoproteins. Gene Ther. 2013 Jan;20(1):16–23. doi: 10.1038/gt.2011.209. Epub 2012 Jan 5. PMID: 22218301.

10. Argaw T, Marino MP, Timmons A, Eldridge L, Takeda K, Li P, Kwilas A, Ou W, Reiser J. *In vivo* targeting of lentiviral vectors pseudotyped with the Tupaia paramyxovirus H glycoprotein bearing a cell-specific ligand. Mol Ther Methods Clin Dev. 2021 Apr 24;21:670–680. doi: 10.1016/j.omtm.2021.04.012. PMID: 34141822; PMCID: PMC8166926.

11. Buchrieser J, Dufloo J, Hubert M, Monel B, Planas D, Rajah MM, Planchais C, Porrot F, Guivel-Benhassine F, Van der Werf S, Casartelli N, Mouquet H, Bruel T, Schwartz O. Syncytia formation by SARS-CoV-2-infected cells. EMBO J. 2020 Dec 1;39(23):e106267. doi: 10.15252/embj.2020106267. Epub 2020 Nov 4. Erratum in: EMBO J. 2021 Feb 1;40(3):e107405. PMID: 33051876; PMCID: PMC7646020.

12. Braga L, Ali H, Secco I, Chiavacci E, Neves G, Goldhill D, Penn R, Jimenez-Guardeño JM, Ortega-Prieto AM, Bussani R, Cannatà A, Rizzari G, Collesi C, Schneider E, Arosio D, Shah AM, Barclay WS, Malim MH, Burrone J, Giacca M. Drugs that inhibit TMEM16 proteins block SARS-CoV-2 spike-induced syncytia. Nature. 2021 Jun;594(7861):88-93. doi: 10.1038/s41586-021-03491-6. Epub 2021 Apr 7. PMID: 33827113; PMCID: PMC7611055.

13. Zhang Z, Zheng Y, Niu Z, Zhang B, Wang C, Yao X, Peng H, Franca DN, Wang Y, Zhu Y, Su Y, Tang M, Jiang X, Ren H, He M, Wang Y, Gao L, Zhao P, Shi H, Chen Z, Wang X, Piacentini M, Bian X, Melino G, Liu L, Huang H, Sun Q. SARS-CoV-2 spike protein dictates syncytium-mediated lymphocyte elimination. Cell Death Differ. 2021 Sep;28(9):2765–2777. doi: 10.1038/s41418-021-00782-3. Epub 2021 Apr 20. PMID: 33879858; PMCID: PMC8056997.

14. Lin L, Li Q, Wang Y, Shi Y. Syncytia formation during SARS-CoV-2 lung infection: a disastrous unity to eliminate lymphocytes. Cell Death Differ. 2021 Jun;28(6):2019–2021. doi: 10.1038/s41418-021-00795-y. Epub 2021 May 12. PMID: 33981020; PMCID: PMC8114657.

15. Rajah MM, Hubert M, Bishop E, Saunders N, Robinot R, Grzelak L, Planas D, Dufloo J, Gellenoncourt S, Bongers A, Zivaljic M, Planchais C, Guivel-Benhassine F, Porrot F, Mouquet H, Chakrabarti LA, Buchrieser J, Schwartz O. SARS-CoV-2 Alpha, Beta, and Delta variants display enhanced Spike-mediated syncytia formation. EMBO J. 2021 Dec 15;40(24):e108944. doi: 10.15252/embj.2021108944. Epub 2021 Oct 25. PMID: 34601723; PMCID: PMC8646911.

16. Rajah MM, Bernier A, Buchrieser J, Schwartz O. The Mechanism and Consequences of SARS-CoV-2 Spike-Mediated Fusion and Syncytia Formation. J Mol Biol. 2022 Mar 30;434(6):167280. doi: 10.1016/j.jmb.2021.167280. Epub 2021 Oct 1. PMID: 34606831; PMCID: PMC8485708.

17. Zeng C, Evans JP, King T, Zheng YM, Oltz EM, Whelan SPJ, Saif LJ, Peeples ME, Liu SL. SARS-CoV-2 spreads through cell-to-cell transmission. Proc Natl Acad Sci U S A. 2022 Jan 4;119(1):e2111400119. doi: 10.1073/pnas.2111400119. PMID: 34937699; PMCID: PMC8740724.

18. Chaudhary S, Yadav RP, Kumar S, Yadav SC. Ultrastructural study confirms the formation of single and heterotypic syncytial cells in bronchoalveolar fluids of COVID-19 patients. Virol J. 2023 May 19;20(1):97. doi: 10.1186/s12985-023-02062-7. PMID: 37208729; PMCID: PMC10198030.

19. Banerjee AK, Blanco MR, Bruce EA, Honson DD, Chen LM, Chow A, Bhat P, Ollikainen N, Quinodoz SA, Loney C, Thai J, Miller ZD, Lin AE, Schmidt MM, Stewart DG, Goldfarb D, De Lorenzo G, Rihn SJ, Voorhees RM, Botten JW, Majumdar D, Guttman M. SARS-CoV-2 Disrupts Splicing, Translation, and Protein Trafficking to Suppress Host Defenses. Cell. 2020 Nov 25;183(5):1325–1339.e21. doi: 10.1016/j.cell.2020.10.004. Epub 2020 Oct 8. PMID: 33080218; PMCID: PMC7543886.

20. Finkel Y, Gluck A, Nachshon A, Winkler R, Fisher T, Rozman B, Mizrahi O, Lubelsky Y, Zuckerman B, Slobodin B, Yahalom-Ronen Y, Tamir H, Ulitsky I, Israely T, Paran N, Schwartz M, Stern-Ginossar N. SARS-CoV-2 uses a multipronged strategy to impede host protein synthesis. Nature. 2021 Jun;594(7862):240-245. doi: 10.1038/s41586-021-03610-3. Epub 2021 May 12. PMID: 33979833.

21. Vora SM, Fontana P, Mao T, Leger V, Zhang Y, Fu TM, Lieberman J, Gehrke L, Shi M, Wang L, Iwasaki A, Wu H. Targeting stem-loop 1 of the SARS-CoV-2 5’ UTR to suppress viral translation and Nsp1 evasion. Proc Natl Acad Sci U S A. 2022 Mar 1;119(9):e2117198119. doi: 10.1073/pnas.2117198119. PMID: 35149555; PMCID: PMC8892331.

22. Hanke L, Vidakovics Perez L, Sheward DJ, Das H, Schulte T, Moliner-Morro A, Corcoran M, Achour A, Karlsson Hedestam GB, Hällberg BM, Murrell B, McInerney GM. An alpaca nanobody neutralizes SARS-CoV-2 by blocking receptor interaction. Nat Commun. 2020 Sep 4;11(1):4420. doi: 10.1038/s41467-020-18174-5. PMID: 32887876; PMCID: PMC7473855.

23. Huo J, Le Bas A, Ruza RR, Duyvesteyn HME, Mikolajek H, Malinauskas T, Tan TK, Rijal P, Dumoux M, Ward PN, Ren J, Zhou D, Harrison PJ, Weckener M, Clare DK, Vogirala VK, Radecke J, Moynié L, Zhao Y, Gilbert-Jaramillo J, Knight ML, Tree JA, Buttigieg KR, Coombes N, Elmore MJ, Carroll MW, Carrique L, Shah PNM, James W, Townsend AR, Stuart DI, Owens RJ, Naismith JH. Neutralizing nanobodies bind SARS-CoV-2 spike RBD and block interaction with ACE2. Nat Struct Mol Biol. 2020 Sep;27(9):846–854. doi: 10.1038/s41594-020-0469-6. Epub 2020 Jul 13. Erratum in: Nat Struct Mol Biol. 2020 Nov;27(11):1094. Erratum in: Nat Struct Mol Biol. 2021 Mar;28(3):326. PMID: 32661423.

24. Karoyan P, Vieillard V, Gómez-Morales L, Odile E, Guihot A, Luyt CE, Denis A, Grondin P, Lequin O. Human ACE2 peptide-mimics block SARS-CoV-2 pulmonary cells infection. Commun Biol. 2021 Feb 12;4(1):197. doi: 10.1038/s42003-021-01736-8. PMID: 33580154; PMCID: PMC7881012.

25. Zhang H, Lv P, Jiang J, Liu Y, Yan R, Shu S, Hu B, Xiao H, Cai K, Yuan S, Li Y. Advances in developing ACE2 derivatives against SARS-CoV-2. Lancet Microbe. 2023 May;4(5):e369–e378. doi: 10.1016/S2666-5247(23)00011-3. Epub 2023 Mar 16. PMID: 36934742; PMCID: PMC10019897.

26. Burns JC, Friedmann T, Driever W, Burrascano M, Yee JK. Vesicular stomatitis virus G glycoprotein pseudotyped retroviral vectors: concentration to very high titer and efficient gene transfer into mammalian and nonmammalian cells. Proc Natl Acad Sci U S A. 1993 Sep 1;90(17):8033–7. doi: 10.1073/pnas.90.17.8033. PMID: 8396259; PMCID: PMC47282.

27. Gutierrez-Guerrero A, Cosset FL, Verhoeyen E. Lentiviral Vector Pseudotypes: Precious Tools to Improve Gene Modification of Hematopoietic Cells for Research and Gene Therapy. Viruses. 2020 Sep 11;12(9):1016. doi: 10.3390/v12091016. PMID: 32933033; PMCID: PMC7551254.

28. Nikolic J, Belot L, Raux H, Legrand P, Gaudin Y, A Albertini A. Structural basis for the recognition of LDL-receptor family members by VSV glycoprotein. Nat Commun. 2018 Mar 12;9(1):1029. doi: 10.1038/s41467-018-03432-4. PMID: 29531262; PMCID: PMC5847621.

29. Gao Y, Bergman I. Vesicular Stomatitis Virus (VSV) G Glycoprotein Can Be Modified to Create a Her2/Neu-Targeted VSV That Eliminates Large Implanted Mammary Tumors. J Virol. 2023 Jun 29;97(6):e0037223. doi: 10.1128/jvi.00372-23. Epub 2023 May 18. PMID: 37199666; PMCID: PMC10308914.

30. Kozlova AA, Verkhovskii RA, Ermakov AV, Bratashov DN. Changes in Autofluorescence Level of Live and Dead Cells for Mouse Cell Lines. J Fluoresc. 2020 Dec;30(6):1483–1489. doi: 10.1007/s10895-020-02611-1. Epub 2020 Sep 1. PMID: 32870453.

31. Bertolo A, Guerrero J, Stoyanov J. Autofluorescence-based sorting removes senescent cells from mesenchymal stromal cell cultures. Sci Rep. 2020 Nov 5;10(1):19084. doi: 10.1038/s41598-020-76202-2. PMID: 33154552; PMCID: PMC7645702.

32. Mace MC, Hansen M, Whiting S, Wang CT, Barklis E. Retroviral envelope protein fusions to secreted and membrane markers. Virology. 1992 Jun;188(2):869–74. doi: 10.1016/0042-6822(92)90544-y. PMID: 1585654.

33. Batéjat C, Grassin Q, Manuguerra JC, Leclercq I. Heat inactivation of the severe acute respiratory syndrome coronavirus 2. J Biosaf Biosecur. 2021 Jun;3(1):1–3. doi: 10.1016/j.jobb.2020.12.001. Epub 2021 Jan 23. PMID: 33521591; PMCID: PMC7825878.

34. Ternette N, Stefanou D, Kuate S, Uberla K, Grunwald T. Expression of RNA virus proteins by RNA polymerase II dependent expression plasmids is hindered at multiple steps. Virol J. 2007 Jun 5;4:51. doi: 10.1186/1743-422X-4-51. PMID: 17550613; PMCID: PMC1892776.

35. Le Hir H, Nott A, Moore MJ. How introns influence and enhance eukaryotic gene expression. Trends Biochem Sci. 2003 Apr;28(4):215–20. doi: 10.1016/S0968-0004(03)00052-5. PMID: 12713906.

36. Rose AB. Introns as Gene Regulators: A Brick on the Accelerator. Front Genet. 2019 Feb 7;9:672. doi: 10.3389/fgene.2018.00672. PMID: 30792737; PMCID: PMC6374622.

37. Diamond MS, Kanneganti T-D. Innate Immunity: the first line of defense against SARS-CoV-2. Nat Immunology 2022 Feb; 23(2): 165–176. Published online 2022 Feb 1. doi: 10.1038/s41590-021-01091-0

38. Sievers BL, Cheng MTK, Csiba K, Meng B, Gupta RK. SARS-CoV-2 and innate immunity: the good, the bad, and the “goldilocks”. Cell Mol Immunol. 2024 Feb;21(2):171–183. doi: 10.1038/s41423-023-01104-y. Epub 2023 Nov 20. PMID: 37985854; PMCID: PMC10805730.

39. Pacheco B, Finzi A, Stremlau M, Sodroski J. Adaptation of HIV-1 to cells expressing rhesus monkey TRIM5α. Virology. 2010 Dec 20;408(2):204–12. doi: 10.1016/j.virol.2010.09.019. Epub 2010 Oct 16. PMID: 20956011; PMCID: PMC2975777.

40. Kuroishi A, Bozek K, Shioda T, Nakayama EE. A single amino acid substitution of the human immunodeficiency virus type 1 capsid protein affects viral sensitivity to TRIM5 alpha. Retrovirology. 2010 Jul 7;7:58. doi: 10.1186/1742-4690-7-58. PMID: 20609213; PMCID: PMC2910007.

41. Soll SJ, Wilson SJ, Kutluay SB, Hatziioannou T, Bieniasz PD. Assisted evolution enables HIV-1 to overcome a high TRIM5α-imposed genetic barrier to rhesus macaque tropism. PLoS Pathog. 2013;9(9):e1003667. doi: 10.1371/journal.ppat.1003667. Epub 2013 Sep 26. PMID: 24086139; PMCID: PMC3784476.

42. Fan Y, Marioli M, Zhang K. Analytical characterization of liposomes and other lipid nanoparticles for drug delivery. J Pharm Biomed Anal. 2021 Jan 5;192:113642. doi: 10.1016/j.jpba.2020.113642. Epub 2020 Sep 19. PMID: 33011580.

43. Alfadhli A, Romanaggi C, Barklis RL, Merutka I, Bates TA, Tafesse FG, Barklis E. Capsid-specific nanobody effects on HIV-1 assembly and infectivity. Virology. 2021 Oct;562:19–28. doi: 10.1016/j.virol.2021.07.001. Epub 2021 Jul 5. PMID: 34246112; PMCID: PMC8419096.

44. Syed AM, Taha TY, Tabata T, Chen IP, Ciling A, Khalid MM, Sreekumar B, Chen PY, Hayashi JM, Soczek KM, Ott M, Doudna JA. Rapid assessment of SARS-CoV-2-evolved variants using virus-like particles. Science. 2021 Dec 24;374(6575):1626-1632. doi: 10.1126/science.abl6184. Epub 2021 Nov 4. PMID: 34735219; PMCID: PMC9005165.

45. Zuffery, R., Nagy, D., Mandel, R., Naldini, L., Trono, D., 1997. Multiply attenuated len-tiviral vector achieves efficient gene delivery in vivo. Nat. Biotechnol. 15, 871–875.

46. Campbell EM, Perez O, Melar M, Hope TJ. Labeling HIV-1 virions with two fluorescent proteins allows identification of virions that have productively entered the target cell. Virology. 2007 Apr 10;360(2):286–93. doi: 10.1016/j.virol.2006.10.025. Epub 2006 Nov 22. PMID: 17123568; PMCID: PMC1885464.

47. Berwin B, Barklis E. Retrovirus-mediated insertion of expressed and non-expressed genes at identical chromosomal locations. Nucleic Acids Res. 1993 May 25;21(10):2399–407. doi: 10.1093/nar/21.10.2399. PMID: 8506135; PMCID: PMC309539.

48. Millán JL. Molecular cloning and sequence analysis of human placental alkaline phosphatase. J Biol Chem. 1986 Mar 5;261(7):3112–5. Erratum in: J Biol Chem 1991 Feb 25;266(6):4023. PMID: 3512548.

49. Adachi A, Gendelman HE, Koenig S, Folks T, Willey R, Rabson A, Martin MA. Production of acquired immunodeficiency syndrome-associated retrovirus in human and nonhuman cells transfected with an infectious molecular clone. J Virol. 1986 Aug;59(2):284–91. doi: 10.1128/JVI.59.2.284-291.1986. PMID: 3016298; PMCID: PMC253077.

50. DuBridge RB, Tang P, Hsia HC, Leong PM, Miller JH, Calos MP. Analysis of mutation in human cells by using an Epstein-Barr virus shuttle system. Mol Cell Biol. 1987 Jan;7(1):379–87. doi: 10.1128/mcb.7.1.379-387.1987. PMID: 3031469; PMCID: PMC365079.

51. Earley EM, Johnson KMThe lineage of Vero, Vero 76 and its clone C1008 in the United StatesIn: Earley EM, Johnson KMVero cells: origin, properties and biomedical applicationsTokyoChiba Univ.pp. 26–29, 1988

52. Longo PA, Kavran JM, Kim MS, Leahy DJ. Transient mammalian cell transfection with polyethylenimine (PEI). Methods Enzymol. 2013;529:227–40. doi: 10.1016/B978-0-12-418687-3.00018-5. PMID: 24011049; PMCID: PMC4012321.

53. Barklis E, Staubus AO, Mack A, Harper L, Barklis RL, Alfadhli A. Lipid biosensor interactions with wild type and matrix deletion HIV-1 Gag proteins. Virology. 2018 May;518:264–271. doi: 10.1016/j.virol.2018.03.004. Epub 2018 Mar 15. PMID: 29549788; PMCID: PMC5911238.

54. Schneider CA, Rasband WS, Eliceiri KW. NIH Image to ImageJ: 25 years of image analysis. Nat Methods. 2012 Jul;9(7):671–5. doi: 10.1038/nmeth.2089. PMID: 22930834; PMCID: PMC5554542.

55. Jones TA, Blaug G, Hansen M, Barklis E. Assembly of gag-beta-galactosidase proteins into retrovirus particles. J Virol. 1990 May;64(5):2265–79. doi: 10.1128/JVI.64.5.2265-2279.1990. PMID: 2109101; PMCID: PMC249388.

56. Alfadhli A, Mack A, Harper L, Berk S, Ritchie C, Barklis E. Analysis of quinolinequinone reactivity, cytotoxicity, and anti-HIV-1 properties. Bioorg Med Chem. 2016 Nov 1;24(21):5618–5625. doi: 10.1016/j.bmc.2016.09.028. Epub 2016 Sep 12. PMID: 27663546; PMCID: PMC5065790.

57. Barklis E, Alfadhli A, Kyle JE, Bramer LM, Bloodsworth KJ, Barklis RL, Leier HC, Petty RM, Zelnik ID, Metz TO, Futerman AH, Tafesse FG. Ceramide synthase 2 deletion decreases the infectivity of HIV-1. J Biol Chem. 2021 Jan-Jun;296:100340. doi: 10.1016/j.jbc.2021.100340. Epub 2021 Jan 28. PMID: 33515546; PMCID: PMC7949126.

58. Walls AC, Park YJ, Tortorici MA, Wall A, McGuire AT, Veesler D. Structure, Function, and Antigenicity of the SARS-CoV-2 Spike Glycoprotein. Cell. 2020 Apr 16;181(2):281–292.e6. doi: 10.1016/j.cell.2020.02.058. Epub 2020 Mar 9. Erratum in: Cell. 2020 Dec 10;183(6):1735. PMID: 32155444; PMCID: PMC7102599.

59. Li M, Ren Y, Aw ZQ, Chen B, Yang Z, Lei Y, Cheng L, Liang Q, Hong J, Yang Y, Chen J, Wong YH, Wei J, Shan S, Zhang S, Ge J, Wang R, Dong JZ, Chen Y, Shi X, Zhang Q, Zhang Z, Chu JJH, Wang X, Zhang L. Broadly neutralizing and protective nanobodies against SARS-CoV-2 Omicron subvariants BA.1, BA.2, and BA.4/5 and diverse sarbecoviruses. Nat Commun. 2022 Dec 27;13(1):7957. doi: 10.1038/s41467-022-35642-2. PMID: 36575191; PMCID: PMC9792944.

60. Caillat C, Guilligay D, Torralba J, Friedrich N, Nieva JL, Trkola A, Chipot CJ, Dehez FL, Weissenhorn W. Structure of HIV-1 gp41 with its membrane anchors targeted by neutralizing antibodies. Elife. 2021 Apr 19;10:e65005. doi: 10.7554/eLife.65005. PMID: 33871352; PMCID: PMC8084527.

